# Crystal structure of NirF: Insights into its role in heme *d*_1_ biosynthesis

**DOI:** 10.1101/2020.01.13.904656

**Authors:** Thomas Klünemann, Manfred Nimtz, Lothar Jänsch, Gunhild Layer, Wulf Blankenfeldt

**Affiliations:** Structure and Function of Proteins, Helmholtz Centre for Infection Research, Inhoffenstrasse 7, 38124 Braunschweig, Germany; Cellular Proteome Research, Helmholtz Centre for Infection Research, Inhoffenstrasse 7, 38124 Braunschweig, Germany; Institute of Pharmaceutical Sciences, Pharmaceutical Biology, Albert-Ludwigs-Universität Freiburg, 79104 Freiburg, Germany; Institute for Biochemistry, Biotechnology and Bioinformatics, Technische Universität Braunschweig, 38106 Braunschweig, Germany

**Keywords:** denitrification, heme *d*_1_, NirF, tetrapyrrole biosynthesis, x-ray structure

## Abstract

Certain facultative anaerobes such as the opportunistic human pathogen *Pseudomonas aeruginosa* can respire on nitrate, a process generally known as denitrification. This enables denitrifying bacteria to survive in anoxic environments and contributes e.g. to the formation of biofilm, hence increasing difficulties in eradicating *P. aeruginosa* infections. A central step in denitrification is the reduction of nitrite to nitrous oxide by nitrite reductase NirS, an enzyme that requires the unique cofactor heme *d_1_*. While heme *d_1_* biosynthesis is mostly understood, the role of the essential periplasmatic protein NirF in this pathway remains unclear. Here, we have determined crystal structures of NirF and its complex with dihydroheme *d_1_*, the last intermediate of heme *d_1_* biosynthesis. We found that NirF forms a bottom-to-bottom β-propeller homodimer and confirmed this by multi-angle light and small-angle X-ray scattering. The N-termini are immediately neighbored and project away from the core structure, which hints at simultaneous membrane anchoring *via* both N-termini. Further, the complex with dihydroheme *d_1_* allowed us to probe the importance of specific residues in the vicinity of the ligand binding site, revealing residues not required for binding or stability of NirF but essential for denitrification in experiments with complemented mutants of a Δ*nirF* strain of *P. aeruginosa*. Together, these data implicate that NirF possesses a yet unknown enzymatic activity and is not simply a binding protein of heme *d_1_* derivatives.

## Introduction

Denitrification, the respiration on nitrate and nitrite, is a facultative trait that enables some bacterial species to proliferate under anaerobic conditions [1]. Denitrification usually consists of a four-step reductive process, starting from nitrate and leading *via* nitrite to nitric oxide (NO), nitrous oxide (N_2_O) and finally to dinitrogen. Denitrifying bacteria are employed to remove N-oxyanions from wastewater; however, they compete with plants for nutrients in agriculture and even affect the climate, as the gaseous intermediates are active greenhouse gases [1]. Furthermore, denitrification also plays a role in infectious disease as exemplified by the opportunistic human pathogen *Pseudomonas aeruginosa*, which resorts to denitrification to support its proliferation in regions depleted of oxygen by the host’s immune system [2].

During denitrification in *P. aeruginosa*, the reduction of nitrite to nitric oxide is catalyzed by the cytochrome *cd_1_* nitrite reductase NirS [3, 4]. This homodimeric enzyme contains an N-terminal cytochrome *c* domain with a covalently attached heme *c* that serves as an electron entry point and a C-terminal eight-bladed β-propeller domain that non-covalently binds heme *d_1_* to function as the active site during nitrite reduction [5]. While the catalytic mechanism of NirS is well understood, some aspects in the biosynthesis of its cofactor heme *d_1_* are still unclear [6, 4].

Heme *d_1_* is an isobacteriochlorin and features two carbonyl groups at rings A and B as well as an acrylate moiety at ring D [7, 8]. The *nir*-operon of denitrifying bacteria encodes most of the enzymes necessary for heme *d_1_* biosynthesis, starting from the common tetrapyrrole precursor uroporphyrinogen III [9–12]. The first steps of heme *d_1_* biosynthesis are the conversion of uroporphyrinogen III to 12,18-didecarboxysiroheme (DDSH) *via* percorrin-2, sirohydrochlorin and siroheme, successively catalyzed by NirE, CysG (in some organisms) and NirDLGH (Figure 1) [13–16]. The intermediate DDSH is further converted by the Radical SAM enzyme NirJ, which sequentially removes the propionate groups from pyrrole rings A and B [17]. Based on mass spectrometry, a tetrapyrrole with two γ-lactones involving the acetate groups at C2 and C7 has been observed as the final product of this reaction, but these lactones may be artifacts of the extraction procedure and alternatives such as hydroxyl groups or hydrogen replacing the propionate chains were proposed [17]. Importantly, the generation of dihydro-heme *d_1_* (dd1), the last intermediate of heme *d_1_* biosynthesis and substrate of the oxidoreductase NirN, was not observed in experiments with NirJ, suggesting that dd1 may be produced by NirF, the only remaining less well characterized protein encoded in the *nir*-operon [18, 17].

**Figure 1:**
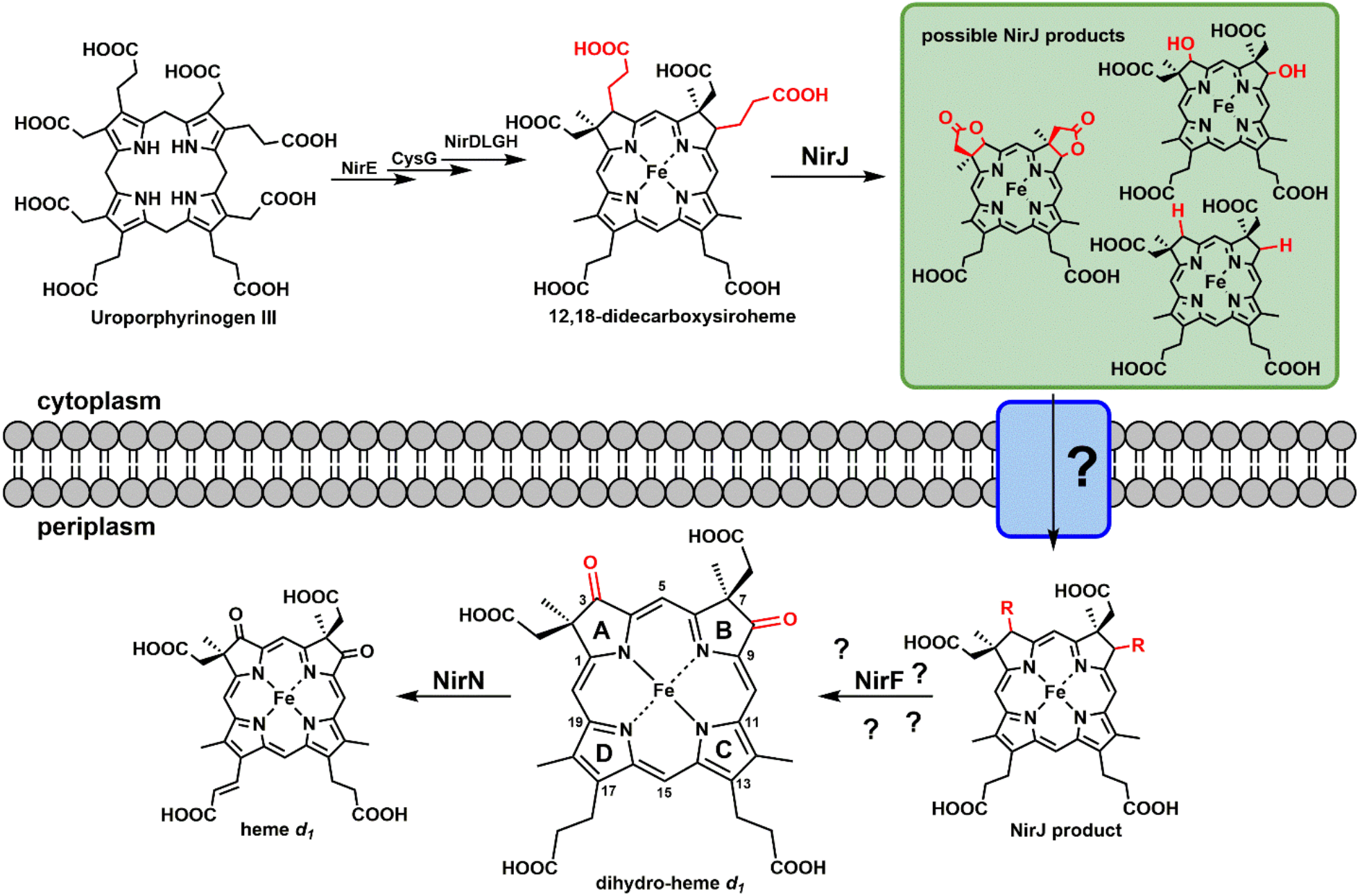
Graphical representation of the heme *d*_1_ biosythesis pathway.

While its exact role still remains unclear, previous work highlighted that NirF is an essential component of the heme *d_1_* biosynthesis pathway [19, 20, 9, 21]. It resides in the periplasm of heme *d_1_*-producing bacteria, where it is even membrane-anchored in some species including *P. aeruginosa*, and this localization was found to be required *in vivo* [21, 22]. It has been demonstrated by UV/Vis spectroscopy that NirF can bind both oxidized and reduced heme *d_1_* as well as dd1 [18, 21]. Exchange of His41 in *Paracoccus pantotrophus* NirF abolished heme *d*_1_ binding, and it was proposed that this residue acts as a proximal iron ligand [21]. Similar to NirN and NirS, NirF is predicted to contain an eight-bladed β-propeller, albeit without being extended by a cytochrome *c* domain. Interestingly, *in vivo* crosslinking experiments in *P. aeruginosa* identified NirN and NirS as transient interaction partners of NirF, hinting at potential substrate shuttling between these proteins [22]. Further, NirF seems to be part of a larger protein complex involved in denitrification, as it was also isolated together with nitric oxide reductase (NorB, NorC) and the electron transfer protein NosR after *in vivo* crosslinking [23]. Together, these data indicate that NirF is an important component of the denitrification process, most likely acting downstream of NirJ and upstream of NirN. To gain further insights into its role, we have determined crystal structures of NirF from *P. aeruginosa* with and without bound dd1 and used these structures to design mutagenesis experiments that highlight the importance of residues in the vicinity of the dd1 rings A and B for the function of the enzyme. Our experiments render further support to the hypothesis that NirF acts on the product of NirJ and provides dd1 to NirN.

## Results and Discussion

### Protein production, purification and crystallization

Heterologous production of *P. aeruginosa* NirF in *E. coli* was established with a construct that lacked the signal sequence for the export into the periplasm and the cysteine residue necessary for membrane attachment via a lipid anchor. By adding N-terminal solubility tags such as SUMO or MBP the amount of recombinant protein could be enhanced up to >500 mg/liter culture. Interestingly, the solubility of purified NirF after tag removal was highly temperature-dependent: when a NirF solution at 8 mg/ml was stored overnight at 4°C, most of the protein precipitated, whereas it was stable when stored at room temperature. Crystallization by the sitting drop vapor diffusion method yielded crystals belonging to the P2_1_ space group containing eight molecules in the asymmetric unit, but analysis with Xtriage [24] revealed severe anisotropy and a large off-origin peak in the Patterson map, indicating the presence of translational non-crystallographic symmetry (tNCS) (Supplementary Table 1). Diffraction data were therefore treated anisotropically using the STARANISO server [25], giving resolution cut-offs of 1.7 Å, 1.6 Å and 2.1 Å along the axes of the fitted ellipsoid. Despite the presence of tNCS, initial phasing was achieved by molecular replacement with an ensemble of the d1-domain of NirN and NirS (for details see the method section). The refinement converged with a model that contains all except the six N-terminal residues and had R-values of R_work_ = 16.3 % and R_free_ = 20.4 %.

### Overall Structure

NirF folds as an eight bladed β-propeller with four antiparallel β-strands per blade and a counterclockwise propagation when viewed from the top (Figure 2). The first blade consists of the three N-terminal and one C-terminal strands and, thus, represents the closing point of this circular protein. This topology allows a C-terminal η-helix to partially close the bottom of the propeller. All other blades propagate from the in- to the outside. Overall, the structure of NirF is highly similar to the d1-domains reported for NirS and NirN, resulting in low RMSDs of 2.4 Å or 2.1 Å, respectively [5, 26].

**Figure 2:**
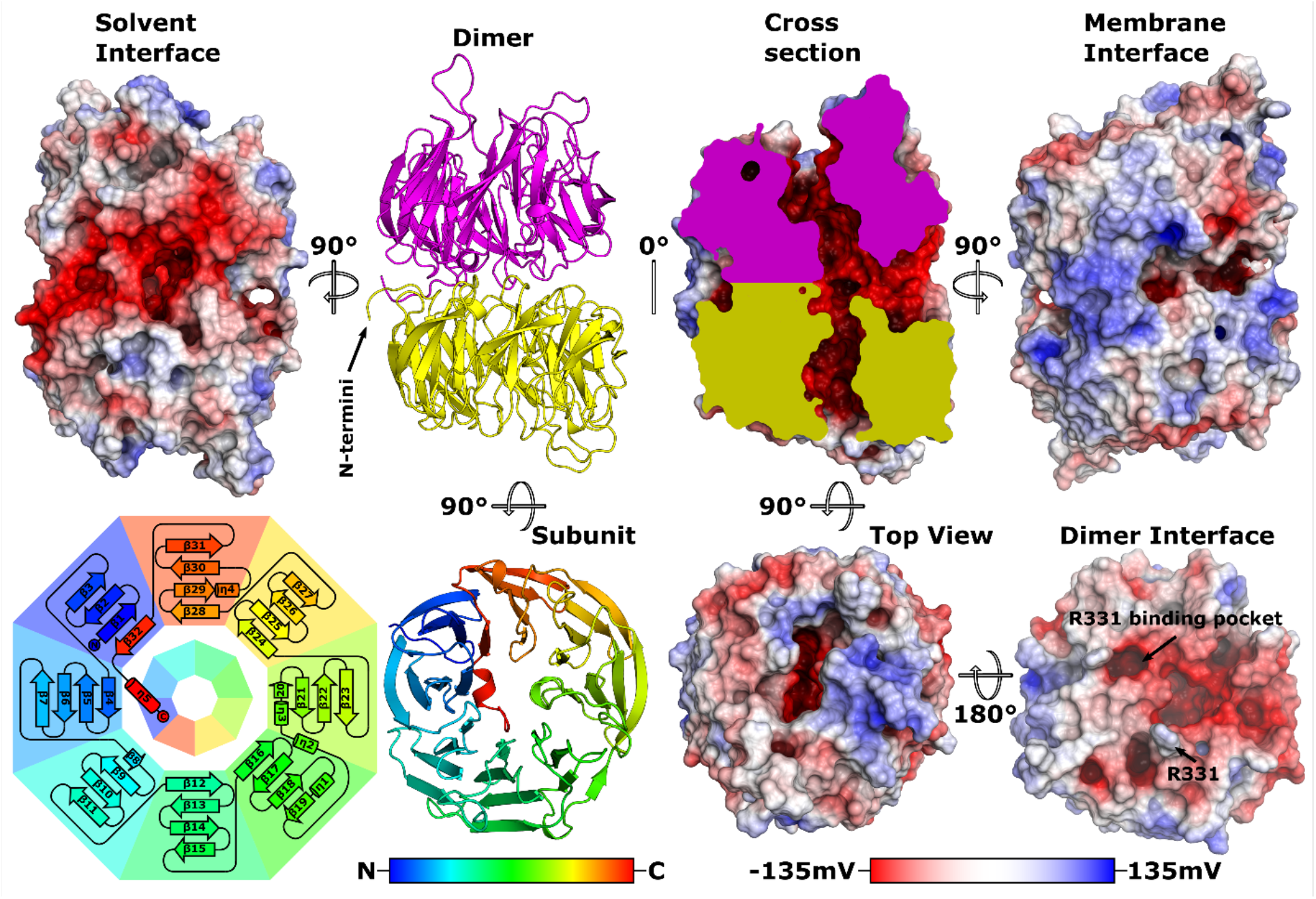
Structure of NirF from *P. aeruginosa* depicted as C_α_ tracing cartoon representation or as a surface model. The surface is colored with a red-white-blue gradient representing the surface potential as calculated by APBS [56]. Different subunits of the dimer are shown in yellow and magenta. Rainbow coloration represents the propagation of the chain. The inner ring of the topology diagram represents the relative orientation of the second subunit. The same color scheme is used throughout.

Analyzing the structure with PISA [27] revealed that NirF has an interface to form homodimers, in which the bottoms of two β-propellers are attached to each other, albeit with low stabilization by solvation free energy (Δ^i^G = − 4 kcal/mol). The two monomers are rotated relative to each other such that the first blade of one propeller is approximately placed between the second and the third of the other (Figure 2 topology diagram). Closer inspection of the interface reveals a crescent-shaped hydrophobic patch that is probably responsible for the negative Δ^i^G values. In addition, the dimers contained in the asymmetric unit are on average stabilized by 15 hydrogen bonds and 22 salt bridges. Of the residues involved in these interactions, R331 seems especially important since it is deeply buried in a negative pocket of the second monomer. Interestingly, a similar bottom-to-bottom dimer of β-propellers has not been reported previously. Proteins of this type usually interact by their sides, which extends their β-sheets, as for example in the d1-domains of NirS [5, 28]. It is possible that the unusual dimerization allows NirF to form side-by-side heterooligomers with the β-propellers of NirN and NirS, which are known transient interaction partners of NirF [22].

To confirm the bottom-to-bottom homodimeric assembly of NirF, we measured small angle x-ray scattering (SAXS) curves. The resulting *ab initio* bead model calculated from these data is consistent with the NirF dimer shown in Fig. 3. Further, theoretical scattering curves and radii of gyration (Rg) derived from other dimer and monomer models of NirF fit the measured data less well, lending additional support to the relevance of the dimer observed *in crystallo*.

**Figure 3:**
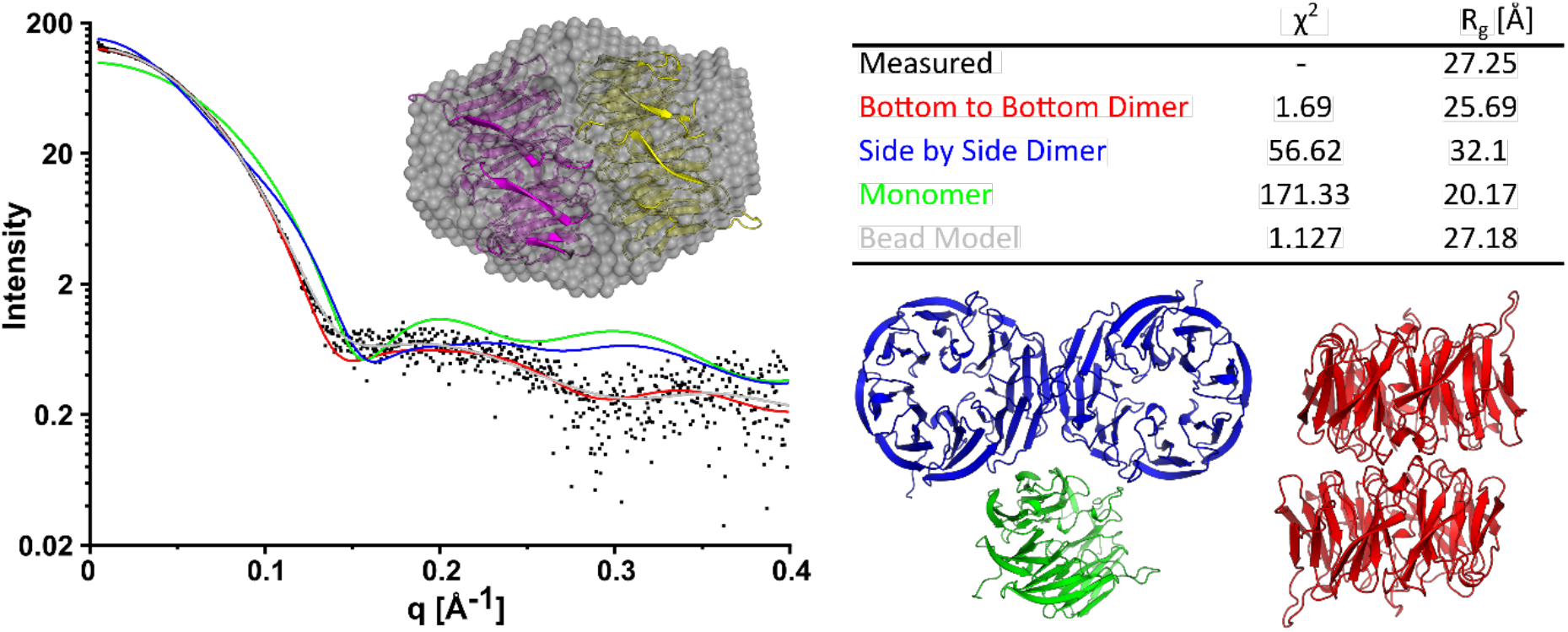
Solution small angle X-ray scattering of NirF of *P. aeruginosa*. Measured data are represented as black dots. Solid lines represent theoretical scattering curves colored according to model used for the calculation. FOXS [55] or Dammin [54] were used for the calculation, χ^2^ values represent goodness of fit to measured data and R_g_ the derived radius of gyration.

An important consequence of the observed dimerization is that the C-terminus must have a defined length, since elongation would interfere with dimerization or tetrapyrrole binding (as shown later). This explains why the NirF variant from *P*a. *pantotrophus*, which was produced with a C-terminal purification tag, was reported to be monomeric and why a *P. aeruginosa* mutant strain lacking NirF but producing a recombinant, C-terminally tagged variant of NirF displayed attenuated growth under denitrifying conditions [22, 21]. Further, twenty-one aligned sequences of NirF from different organisms all end at the same position and with a hydrophobic amino acid (Supplementary Figure 1), indicating that dimerization might be a common feature of NirF.

As NirF of *P. aeruginosa* was found to be membrane-anchored through its N-terminus [22], it is interesting to note that both N-termini are positioned next to each other in the dimer (Figure 2). The surface facing the membrane possesses an overall positive charge, which seems to enable interactions with phospholipids. In contrast, the solvent-exposed side and even more so the interior of the *d_1_*-domains exhibit a mostly negative character. Surprisingly, the insides of both monomers are connected via a narrow T-shaped tunnel that has an additional exit towards the solvent-exposed side. Using MOLEonline 2.0 [29], the radius of the widest bottleneck of this tunnel was determined to be 2 Å, which is too narrow to allow for the passage of tetrapyrroles. Therefore, the tunnel likely represents a solvent channel that allows water to access the tetrapyrrole binding site (next paragraph).

### Dihydro-heme *d_1_* binding site

Crystals of NirF were soaked with dd1 to gain a detailed insight into tetrapyrrole binding. Well-defined |F_O_F_C_| difference electron density with anomalous difference density at the expected position of the iron atom allowed placing dihydro-heme *d_1_* (Figure 4). The model was refined at 2.1, 1.9 and 2.7 Å along the axes of the fitted ellipsoid and gave R-values of R_work_ = 21.5% and R_free_ = 25.8%. Occupancies of the ligand bound to the eight chains contained in the asymmetric unit refined to values between 75% and 100% (details in the method section). Not surprisingly, dd1 is bound in the middle of the d1-domain, similar to other d1-type hemes in NirN and NirS, despite being slightly tilted in the binding pocket [26, 5] (Figure 4 and Supplementary Figure 2). The southern part of the tetrapyrrole is deeply buried in the negatively charged binding pocket, and a positive surface patch accommodates the acetate groups of the northern half. Interestingly, the propionate moiety of ring D is oriented towards the propionate of ring C. This may lead to a discrimination between dihydro-heme *d*_1_ and heme *d*_1_, since the latter carries an acrylate side chain at the D-ring that tends to adopt a planar conformation with respect to the tetrapyrrole ring system, allowing electron delocalization through the π-orbital system as e.g. seen in the structure of NirS [5]. In NirF, such a conformation would lead to clashes with H325.

**Figure 4:**
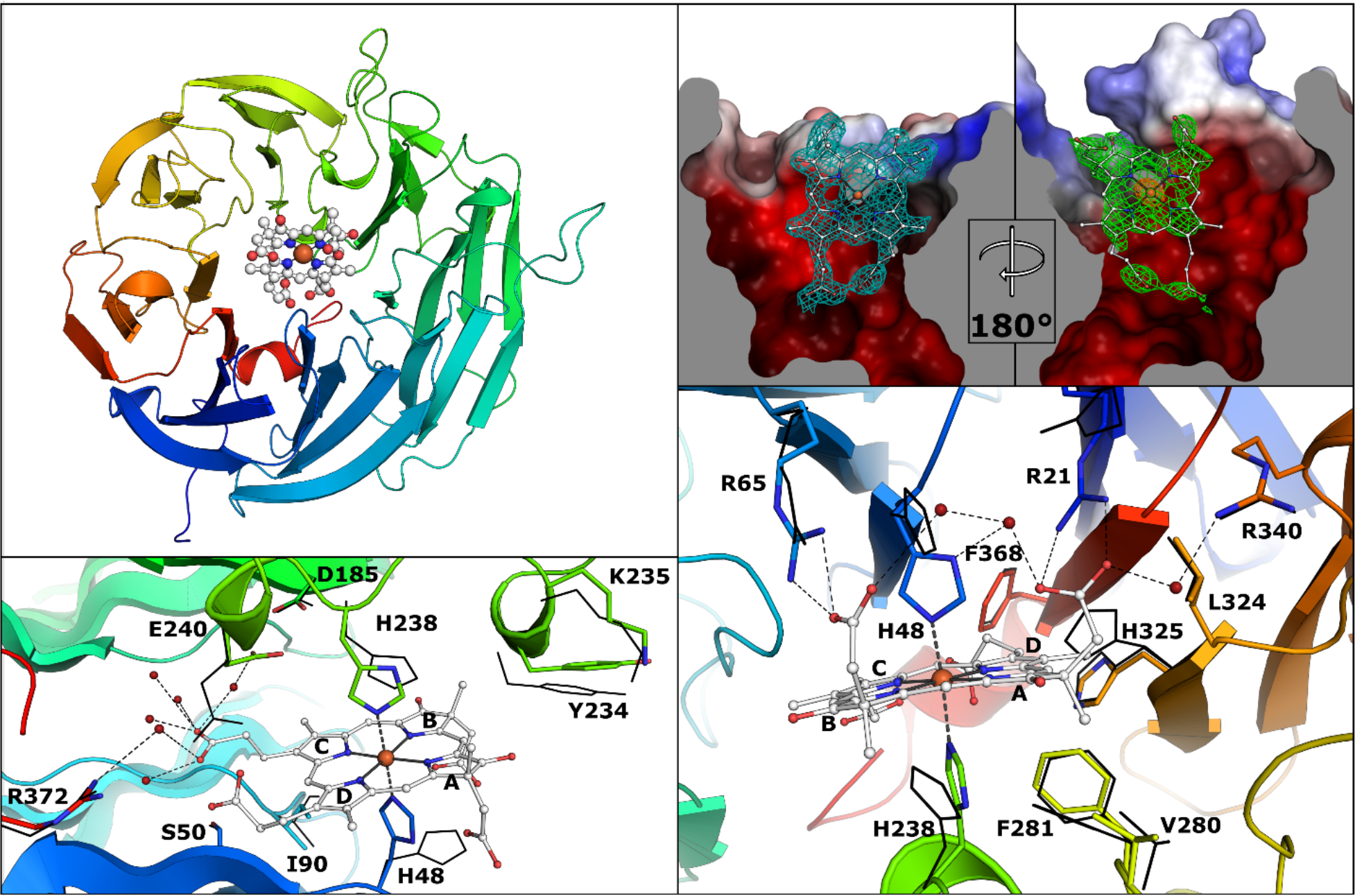
Depiction of the dd1 binding side. The backbone is depicted as cartoon model, residues as sticks, dd1 as white ball-and-stick model and water molecules as red spheres. Positions of residues without bound dd1 are depicted as black lines. Thin dashed black lines represent H-bonds. The cyan mesh represents the final 2F_o_F_C_ map at a σ-level of 1, the green mesh depicts the F_o_F_c_ map prior to first placement of a dd1 molecule at a σ-level of 3 and the orange mesh presents the anoumalous map at a σ-level of 15.

Closer inspection reveals that upon binding of dd1 H48 and H238 rotate to ligate the central iron of the tetrapyrrole (Figure 4). Hydrophobic interaction partners of dd1 are I90, V280, F281, L324 and F368. E240 reorients to avoid clashes with the incoming dd1 and interacts with the propionate of ring C via a water molecule. In addition, the propionate moiety interacts with several water molecules located in the T-shaped tunnel mentioned above, which e.g. establishes a bridge to R372. The acetate groups of the northern half of dd1 form hydrogen bonds with R21 and R65 and, again through bridging water molecules, to R340 and H48. S50 and D185 are found in the vicinity of dd1, but do not seem to contact the ligand. Surprisingly, neither direct nor indirect interactions with the oxo groups of dd1 are present in the final model. A comparison of 21 NirF sequences (seven of each class of α-, β- and γ-proteobacteria) shows that, except for E240 and V280, all of the mentioned residues are identical (Supplementary Figure 1). V280 is sometimes found as an isoleucine, whereas E240 is replaced by glutamine or arginine in some cases.

### Comparison to NirN and NirS

As mentioned above, an eight-bladed β-propeller that binds d1-type hemes is also found in the dehydrogenase NirN and in the nitrite reductase NirS [26, 5]. The unknown but essential role of NirF must differ from these proteins, suggesting that dd1-binding residues that are different from those found in NirN or NirS are involved in its function (Supplementary Figure 2 and 3). The dd1 iron-ligating distal H48 and proximal H238 are conserved in all three proteins, however the proximal histidine of NirS does not coordinate the iron directly, but rather is involved in substrate (nitrite) binding. The hydrophobic nature of V280, F281, L324 and F368 is conserved across the three proteins, whereas the equivalent position of I90 is found as an arginine that interacts with the oxo- or acetate groups of the B-ring in NirN and NirS. Interestingly, the northern half of dd1 interacts with three arginines in all three proteins, which, with the exception of R65, originate from different positions of the β-propeller. This makes the NirF-specific R340 an interesting candidate for further investigation. Other residues that are unique to NirF are S50, E240 and R372. Another overall unique feature of NirF is the loop that connects β-sheets five and six (residues 226-236, loop 1), which is more extended than in NirN and NirS and places Y234 as well as K235 nearly on top of dd1, implying functional involvement of these NirF-conserved residues. Loop 2 (residues 126-139), connecting β-sheets three and four, adopts a different conformation in all three proteins. As this loop is involved in dimerization in NirS, it might fulfill a similar interfacing function in the previously detected transient heterooligomers of NirF with NirN or NirS [22]. We have therefore devised experiments to probe the importance of these amino acids, also including D185, since this residue of the tetrapyrrole binding site is conserved across all three β-propeller proteins, but has not been analyzed to date.

### Impact of amino acid exchanges in NirF on denitrification by *P. aeruginosa*

Because the function of NirF is not known yet, we used the crystal structure in complex with dd1 to design variants of the protein and assess their effect on the denitrification capacity of *P. aeruginosa* to gain new insight into the role of this protein. Toward this, we employed the *P. aeruginosa* PA01 Δ*nirF* strain RM301 [9] carrying plasmids encoding versions of NirF in which residues in the vicinity of the ligand binding site had been replaced by alanine. For these newly generated *P. aeruginosa* strains, growth curves were measured under denitrifying conditions. Additionally, the nitrate consumption and the accumulation of nitrite in the media was determined. This system was already employed to confirm the location of NirF in the periplasm in earlier work [22, 21]. As shown in Fig. 5 and as reported previously [9, 22], *P. aeruginosa* strain RM301 indeed displayed impaired growth under denitrifying conditions, whereas the strain RM301 that produced wild type NirF from a plasmid even outgrew the parent strain.

**Figure 5:**
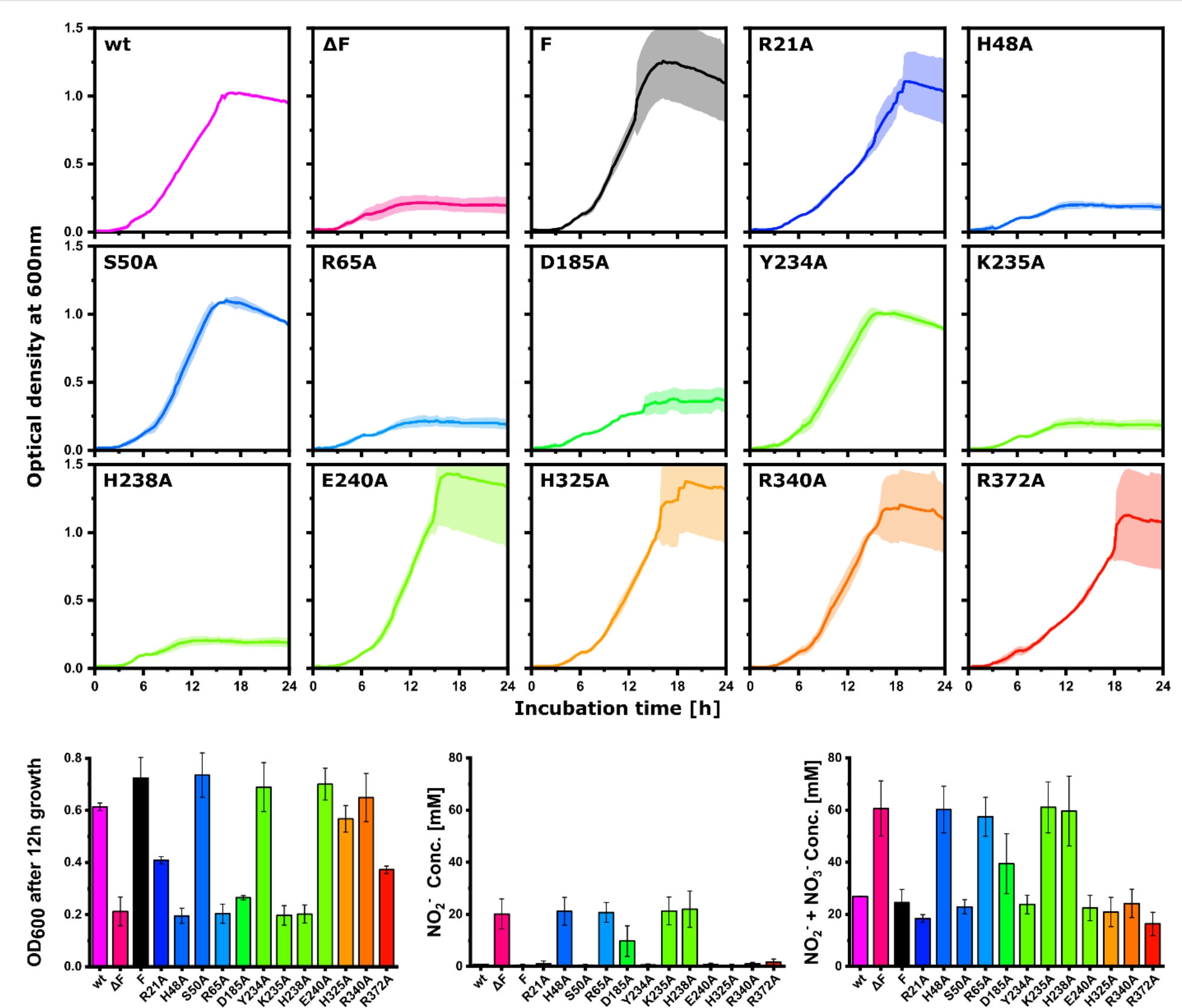
Averaged growth curves of four biological replicates of the *P. aeruginosa* PAO1 strain RM301 complemented with plasmid-born NirF containing different amino acid exchanges (wt only two replicates). Errors are marked as pale broadened lines. Large errors towards the end of the experiment were due to the occurrence of nitrogen gas bubbles, confirming denitrification activity in the culture. Samples marked as wt or ΔF are wild type PA01 or strain RM301 containing only the plasmid backbone, respectively. F designates strain RM301 containing a plasmid with unmutated *nirF*. Bar graphs depict the OD_600_ after 12h taken from the growth curves for easy comparison, and the media concentration of NO_2_^−^ and NO_3_^−^ after 24h of incubation. The coloration correlates to residue positions in the sequence as in shown in Figure 4.

The complementation of strain RM301 with NirF variants carrying exchanges of S50, Y234, E240, H324 and R340 against alanine had no influence on the growth of the respective strain, suggesting that these residues are not required for the activity of NirF. Exchange of R21 or R372, on the other hand, led to attenuated growth but without accumulation of nitrite, hinting at a probably reduced level of active nitrite reductase NirS and hence impaired heme *d*_1_ production. Surprisingly, growth was not only inhibited after exchange of the iron-ligating histidines H48 and H328, but also when the propionate group-coordinating R65 and K235 were replaced by alanine. Finally, exchange of D185 also led to impaired growth, albeit some residual NirS activity seemed to be present.

### Impact of amino acid exchanges on the biophysical properties of NirF

Because the impaired growth of *P. aeruginosa* RM301 producing altered versions of NirF could be rooted in inherent instability of the respective protein variant, we also produced and purified the recombinant proteins and tested for their thermal stability as well as dimerization and dd1 binding (Figure 6 and 7). All of the NirF variants, with exception of the D185A amino acid exchange, could be purified with the procedure established for the wild type protein. The D185A NirF, however, precipitated within minutes after the addition of SUMO-protease to remove the SUMO solubility tag, and only small amounts of the protein could be recovered and purified. Interestingly, the melting point of the purified protein was only 3.1 K lower than that of the wild type protein, which is considerably less than the 13.4 K lower melting temperature for R372A NirF, a variant that could be produced without difficulties. In addition, the D185A variant bound dd1 with similar efficiency as the wild type protein. Together with the observation that denitrification was still active in the respective *P. aeruginosa* strain, this indicates that exchange of D185 impairs the folding but not the stability of the protein and that this residue is probably also not involved in the process purported by NirF.

**Figure 6:**
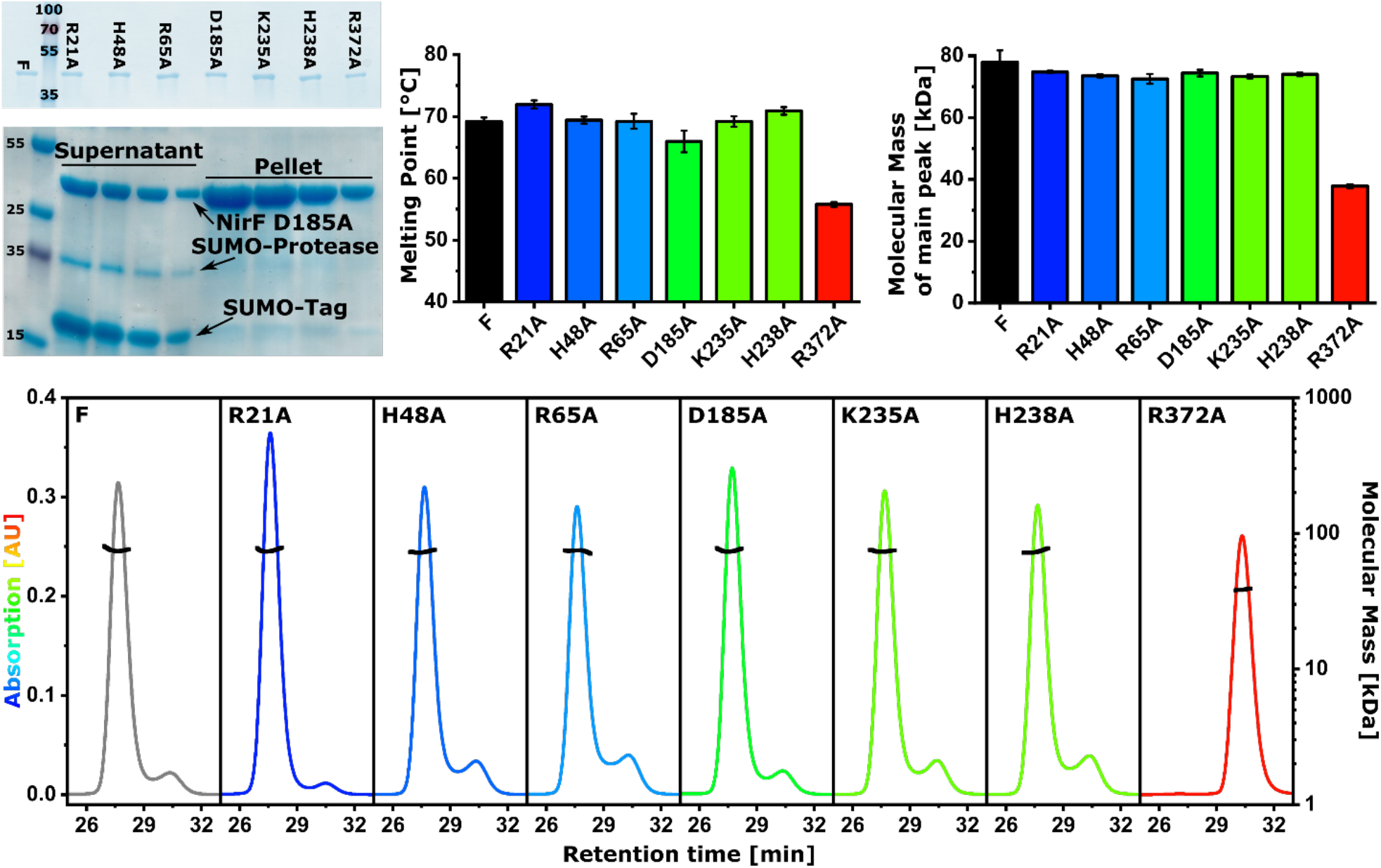
SDS gels of the purified NirF wild type (F) and mutant samples indicate high purity and similar size. The second SDS gel shows different dilutions of supernatant and pellet samples taken from NirF D185A after SUMO protease digestion and centrifugation. Bar diagrams show the melting points and molecular weight of NirF derivates determined by differential scanning fluorometry and SEC-MALS measurements. Error bars represent standard deviations of triplicate measurements. In the bottom panel, representative SEC-MALS chromatograms are depicted with calculated molecular masses shown as black and UV-absorption as colored lines.

**Figure 7:**
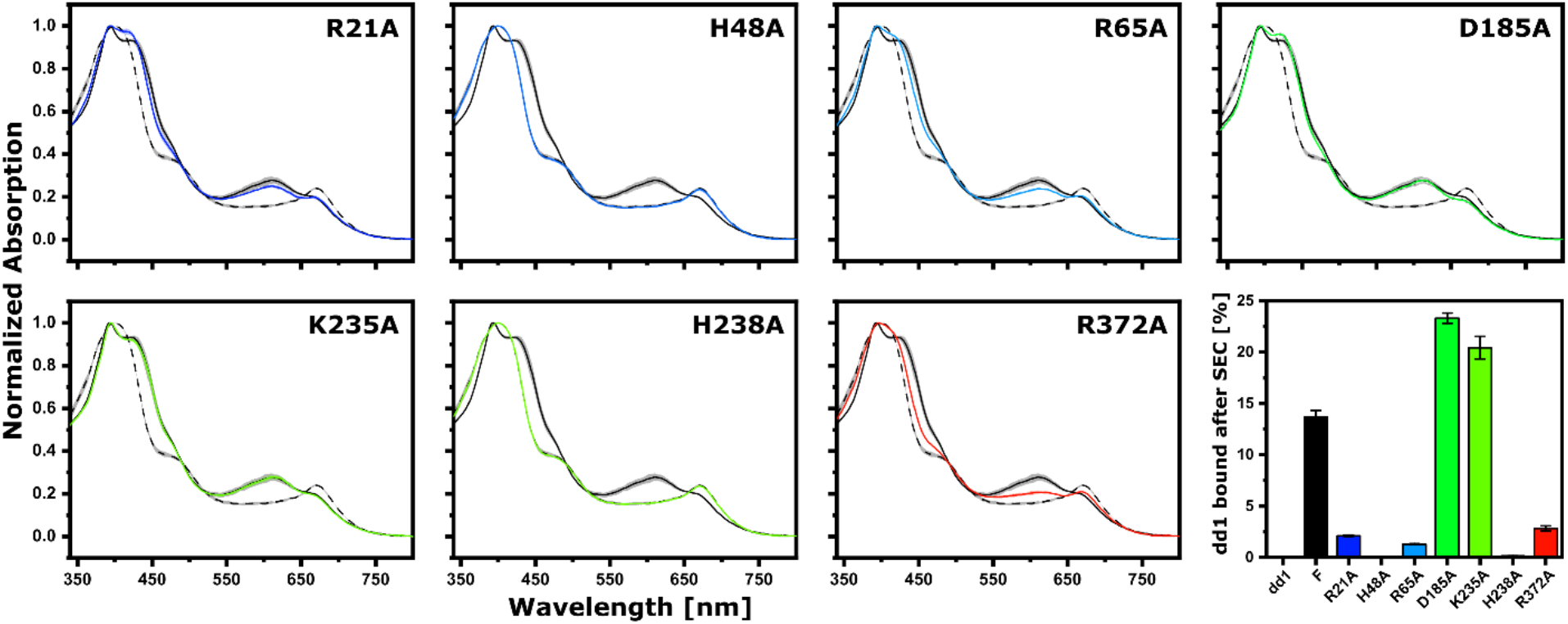
Normalized, averaged and smoothed UV-Vis spectra of dd1 binding to NirF variants. Black dashed lines represent spectra of dd1 not bound to NirF, solid black lines depict spectra of dd1 bound to wild type NirF and colored lines illustrate spectra of dd1 incubated with NirF variants. Gray/Colored shadows represent the standard deviation. The bar diagram depicts the amount of dd1 coeluted with NirF after size exclusion chromatography. Equimolar amounts were injected.

The large decrease in the melting point of the R372A variant without significantly compromising growth under denitrifying conditions prompted us to analyze this variant further. Interestingly, the R372A protein eluted later in size exclusion chromatography and showed a molecular weight corresponding to a NirF monomer in multi-angle light scattering measurements. Although R372 is not involved in the dimer interface, it might stabilize the C-terminal η-helix on which it resides. Changes within this helix might interfere with dimerization, which stabilizes the protein but does not seem to have functional importance, a well-known behavior for many homodimeric proteins [30].

The binding of dd1 to NirF was investigated by UV/Vis absorption spectroscopy after mixing the protein and the tetrapyrrole (Figure 7). While the isolated dd1 exhibits characteristic absorption maxima at 401 nm (Soret band) and 683 nm, the binding of the tetrapyrrole to wild type NirF leads to a split Soret band with maxima at 393 nm and 420 nm as well as a shift of the 683 nm feature to 612 nm. Additionally, dd1 binding to NirF was also assessed by testing for co-elution from a size exclusion column (Figure 7). Both methods demonstrated that the exchange of histidines H48 and H238, which coordinate the central iron atom of dd1, completely abolished terapyrrole binding. Exchange of arginines R21, R65 and R372 resulted in reduced dd1 binding, whereas replacement of D185 and K235 surprisingly increased it. Together, these experiments hint at important functional roles for R65 and K235, since the respective NirF variants are stable and bind dd1 *in vitro*, but are not able to complement the *P. aeruginosa* RM301 strain *in vivo*.

### Conclusions

Previous studies have shown that NirF is required for heme *d*_1_ biosynthesis [9, 21]. Additionally, NirF was found to be periplasmic and, in certain bacteria such as the opportunistic pathogen *P. aeruginosa*, membrane-anchored via its N-terminus [22]. Other components of the denitrification cascade interact with the protein; however, the exact role of NirF is still unclear [23]. The work reported here provides a structural basis for unravelling this role.

Crystal structure analysis revealed that NirF adopts the predicted eight-bladed β-propeller fold and that the protein from *P. aeruginosa* is homodimeric with an unusual bottom-to-bottom setup that has not been observed previously, prompting us to confirm the quarternary structure by SAXS. The arrangement of the dimer positions the N-termini of both monomers in close proximity to each other. Both N-termini point to the same outward-projecting direction with respect to the core structure, in line with simultaneous membrane anchoring through the N-terminus of both subunits. Despite these observations, our experiments with the R372A variant revealed that dimerization is not essential for the unknown function of NirF, which seems to resonate with the observation that the two binding sites for heme *d_1_* derivatives of the dimer locate opposite to the dimerization interface and are structurally independent of each other.

The NirF-dd1 complex structure revealed the position of the tetrapyrrole binding site as well as structural alterations that accompany dd1 binding, which indicate the physiological relevance of the latter. Further, several amino acid residues were identified that may be involved in the function of NirF. These residues were further investigated for their importance in denitrification by *P. aeruginosa in vivo* and in dd1 binding *in vitro*. The most striking effect was observed for histidines H48 and H238, which are absolutely essential for *in vivo* function and *in vitro* tetrapyrrole binding. Again, this finding supports the physiological significance of dd1 binding to NirF. In this context, it is interesting to note that dd1 is relatively solvent exposed and that its A- and B-rings point outwards in the NirF complex. This leads us to speculate about two alternative scenarios for the role of the protein. First, as has been suggested previously, NirF might be responsible for the uptake of dihydro-heme *d*_1_ after its translocation through the membrane and before further transfer to NirN, which catalyzes the last step of heme *d*_1_ biosynthesis [18, 22]. However, in this scenario it is not obvious why NirF is absolutely essential for heme *d*_1_ formation. Alternatively, a role of NirF as a biosynthetic enzyme could be envisaged, potentially involved in the introduction of the carbonyl functions at C3 and C8 of rings A and B. While it has been shown that the cytoplasmic Radical SAM enzyme NirJ removes the two propionate moieties at C3 and C8 of 12,18-didecarboxysiroheme (DDSH) [17] and that the periplasmic NirN converts the propionate at C17 to an acrylate group in the final step of the pathway [18, 17], the formation of the carbonyl functions at C3 and C8 is still enigmatic. Although it was speculated that NirJ might also be responsible for the oxidation of these position, the installation of the carbonyls by NirJ has not been observed experimentally so far [17, 31]. Therefore, we hypothesize that the oxidation at C3 and C8 might be conducted by NirF. However, NirF does not possess a binding site for familiar redox cofactors such as NAD^+^ or FAD [21] and would therefore require an alternative electron acceptor for enzymatic turnover. In this regard, it is interesting to note that the dehydrogenation reaction catalyzed by the structurally related NirN does also not rely on NAD^+^ or FAD as redox cofactors, but instead uses the cytochrome *c* moiety of the enzyme itself as well as the reaction product as the electron acceptors. Since NirF does not possess a cytochrome *c* domain, an external redox-active protein could serve as such an electron acceptor. For electron transfer, this acceptor would possibly have to dock to NirF, and the outward-facing K235 might be involved in this interaction, providing a possible explanation for its requirement for denitrification in the cell-based experiments described above. The *nir*-operon of *P. aeruginosa* encodes two *c*-type cytochromes, NirC and NirM, and the role of NirC is not fully understood. Thus, NirC might act as a redox partner of NirF in *P. aeruginosa*. We are currently devising experiments to test this hypothesis.

## Methods

### Chemicals

All chemicals and media ingredients were purchased from Sigma-Aldrich (St. Louis, U.S.A.) or from Carl Roth GmbH (Karlsruhe, Germany) unless stated otherwise.

### Bacteria and plasmids

The bacterial strains and plasmids used in this study are listed in supplementary Table 2 and 3. The *P. aeruginosa* strains PA01 RM301 (*nirF∷tet*) [9] and RM361 (*nirN∷tet*) [10] were a kind gift by Dr. Hiroyuki Arai. The plasmid pUCP20T-*nirF* was from a previous study [22]. pMALX-E was provided by the Pedersen Lab [32], pET-SUMO is a derivative of pET19 (Invitrogen).

### Construction of Vectors

pUCP20T-*nirF* was used to PCR-amplify *nirF* without the N-terminal periplasmic export signal and the cysteine required for membrane anchoring. pET-SUMO and pMALX-E were digested with Nde1/BamHI or Nhe1/ HindIII (New England Biolabs, Frankfurt, Germany), respectively. The PCR products were ligated with a QuickFusion kit (Absource Diagnostics, Munich, Germany) and *E. coli* XL1-blue as a cloning host, resulting in plasmids pET-SUMO-*nirF* and pMALX-E-*nirF* encoding for the truncated *nirF* gene with an N-terminal 6xHis-SUMO- or MBP-tag. Site directed mutagenesis of pUCP20T-*nirF* and pET-SUMO-*nirF* was performed by QuikChange mutagenesis. Primers used in this study are listed in Supplementary Table 4. Prior to use, all plasmids were checked by Sanger sequencing using services provided by Eurofins (München, Germany) or SeqLab (Göttingen, Germany).

### Transformation and bacterial cultures

Rubidium-competent *E. coli* cells were transformed by heat-shock and selected on LB-agar (BD Bioscience, Franklin Lakes, U.S.A.) plates supplemented with 100 μg/ml ampicillin [33]. *P. aeruginosa* strains were transformed by electroporation using a GenePulser II (BioRad, Hercules (CA), United States) and selected on LB-Agar plates supplemented with 250 μg/ml carbenicillin [34]. Overnight cultures were started by inoculating LB-media containing the required antibiotics with single colonies. The production of SUMO-NirF and MALX-E-NirF in *E. coli* BL21 was performed in 1 L auto induction media [35]. Cultures were incubated for 4 h at 37 °C before the temperature was reduced to 20 °C and growth was continued for another 12 to 16 h. For the production of dd1, *P. aeruginosa* PA01 RM361 was grown under denitrifying conditions as previously described [18, 26]. Cells were harvested by centrifugation, frozen in liquid nitrogen and stored at −20 °C until needed.

### *Pseudomonas* growth curves and nitrite/nitrate determination

Aerobically grown overnight cultures of *P. aeruginosa* were used to inoculate a modified AB minimal medium [36] supplemented with 75 mM NaNO_3_, 200 μM 5-aminolevulinic acid (Alfa Aesar, Haverhill,U.S.A), 200μM Fe(II)SO_4_ and 250μM carbenicillin to an optical density of approximately 0.005. 200 μl of culture were filled in Honeycomb plates and overlaid with 200 μl of paraffin oil to prevent O_2_ exchange with the air. Growth at 37°C was monitored in a Bioscreen C instrument (Oy Growth Curves Ab Ltd, Helsinki, Finland). After 24 h the cultures were harvested and centrifuged in a table top centrifuge.

Nitrite and nitrate concentrations of the supernatant were determined using a colorimetric assay kit (Cayman chemicals, Ann Arbor, United States). For each strain, growth curves of four biological replicates with three technical replicates for each were collected. For some technical replicates, bubbles started to accumulate after ~12 h between oil and culture, leading to discontinuous growth curves. These cultures were excluded from the analysis, if the effect already occurred before the 12 h mark. The remaining technical replicates were averaged to give the growth curve of one biological replicate. The average of all four biological replicates was then produced for the final analysis.

### Purification of recombinant NirF

Purification of MALX-E-NirF and SUMO-NirF was conducted in a buffer consisting of 10 mM Tris pH 8.0, 500 mM NaCl and 10 % (v/v) glycerol (protein buffer). *E. coli* cell pellets were resuspended in protein buffer containing cOmplete protease inhibitor cocktail (Roche, Basel, Switzerland), lysed by sonication and centrifuged for 45 min at 38,000 *g* to remove cell debris. All further steps were performed at room temperature since NirF derivatives were found to be less soluble at low temperature (see main text). To purify SUMO-NirF and its variants, the cleared lysate was loaded onto a 5 ml HisTrap FF column (GE Healthcare, Chicago, U.S.A.) connected to an Äkta Purifier FPLC system (GE Healthcare, Chicago, U.S.A.). The column was washed with three column volumes of protein buffer supplemented with 20 mM imidazol before elution with a gradient to 250 mM imidazole. The SUMO-tag was then removed with 6x-His-tagged SUMO-protease in the course of overnight dialysis against protein buffer containing 1 mM of DTT. To separate the cleaved tag and the protease, NirF was again loaded onto the 5 mL HisTrap FF column and eluted with a gradient to 50 mM imidazole. MALX-E-NirF was purified by loading the cleared cell lysate onto a 5ml MBPtrap HP column (GE Healthcare, Chicago, United States) which was eluted with a gradient to 20 mM maltose. The MBP-tag was not removed for subsequent experiments. All proteins were further purified by size exclusion chromatography using a HiLoad 26/60 Superdex200 pg column (GE Healthcare, Chicago, United States). The purity of proteins was confirmed with SDS-PAGE and concentrations were determined photometrically, using a NanoDrop 2000 (ThermoFisher Scientific, Waltham, U.S.A.) with extinction coefficients calculated by Protparam [37]. The identity of the isolated proteins and presence of assigned amino acid exchanges were checked by mass spectrometry.

### Purification of dihydroheme *d*_1_

Dihydroheme *d*_1_ was obtained by solvent extraction from purified NirS from *P. aeruginosa* PA01 RM361 using acidified ethyl acetate as described by Adamczack *et al*. [18]. Concentrations of dd1 in solution were determined with extinction coefficients published previously [26].

### Crystallization of NirF

Crystallization experiments were set up with a HoneyBee 961 robot (Digilab Genomic Solutions, Hopkinton, U.S.A.) mixing 200 nL of reservoir with 200 nL of protein solution equilibrated against 60 μL of reservoir in 96-well sitting drop vapor diffusion plates (Intelli 96-3 plates, Art Robbins Instruments, Sunnyvale, U.S.A.). Initial crystallization conditions were identified with commercial sparse matrix screens JCSG+ (Qiagen, Hilden, Germany), Index (Hampton Research, U.S.A.), Morpheus (Molecular Dimension, Newmarket, UK) and PEG Suite (Qiagen, Hilden, Germany). Diffracting crystals of the MALX-E-NirF fusion protein were obtained with 30 mg/ml protein and a precipitant containing 20% (w/v) polyethylene glycol 3350 and 0.2 M ammonium formate. Crystals of free NirF, purified from the SUMO-NirF fusion protein, were obtained with several precipitants, but only those containing PEG 3350 or PEG 4000 gave crystals that diffracted x-rays. Optimizing these conditions yielded crystals of similar quality but never surpassed one directly fished from the Morpheus screen. Presented data are from crystals grown in Morpheus condition C7 and H7 as described by Gorrec *et a*l. [38] using protein concentration of 5 mg/mL or 10 mg/mL. A complex with dd1 was obtained by soaking in ligand-saturated mother liquor for one day. Cryoprotection was achieved by washing in reservoir solution containing 10% of (*R*,*R*)-2,3-butandiol (Alfa Aesar) for a few minutes prior to flash-cooling in liquid nitrogen.

### Data Collection and Processing

2000-3600 diffraction images per crystal were collected at the beamline P11 at PETRA III (DESY, Hamburg, Germany) [39] on a PILATUS 6M fast detector with an oscillation angle of 0.1° per images. Reflections were indexed and integrated with DIALS [40]. Integrated reflections were further processed with POINTLESS [41] and AIMLESS [42]. As the datasets of NirF crystals showed strong anisotropy, they were truncated with a local I/σ(I) of 1.2 and corrected using the STARANISO server [25]. Quality of data was assessed using Xtriage confirming the presence of translational non-crystallographic symmetry in all datasets [24]. Processing and data quality statistics can be found in supplementary table 1.

### Structure determination and refinement

Initial phases were first obtained for the low-resolution diffraction data of the MALX-E-NirF fusion protein, using molecular replacement in PHASER [43]. Search models consisted of a crystal structure of MBP (PDB: 4QVH [44]) and an ensemble of the heme *d*_1_ binding domains of NirN (PDB:6RTE [26]) and NirS (PDB:1NNO [45]). Based on the resulting electron density the search model for the heme *d*_1_ binding domains was truncated containing to approx. 80% of all residues. This modified model was then sufficient for molecular replacement in the diffraction data obtained from crystals of the untagged protein. Structures were subsequently refined by manual adjustments in COOT [46] and computational optimization in REFMAC5 [47] of the CCP4 suite [48] and phenix.refine [49] of the phenix suite [50], applying TLS refinement [51] in the last steps of the procedure. Dihydroheme *d*_1_ was placed in the |F_O_-F_C_|difference electron density appearing if the crystals were soaked. For a first estimate of the ligand occupancy, an anomalous map of the iron signal was calculated with ANODE [52] first. After refinement with fixed occupancies, the average B-factors of the ligand were compared to those of the surrounding protein chains and the occupancies were then adjusted in 0.05 steps until the average B-factors were similar, resulting in occupancies between 75% and 100%. Depictions of the models were made using the PyMol molecular graphics system (Schrödinger LLC, version 2.3.2). Model and refinement statistics are summarized in Supplementary Table 1. The structures can be accessed under the PDB codes #### and ####.

### Small-angle x-ray scattering (SAXS)

Small angle X-ray scattering experiments were performed at beamline BM29 BioSAXS of the European Synchrotron Radiation Facility (ESRF, Grenoble, France) [53] on a Pilatus 1M detector. Scattering data were measured online after 150 μg or 450 μg NirF were separated on a 10/300 Superdex 200 increase size exclusion column utilizing protein buffer without glycerol. Data processing and bead model calculation were performed with the ATSAS software package (version 2.8.4 [54]). Theoretical SAXS profiles of PDB models were calculated with the FoXS server [55]. The side-by-side homodimer model of NirF shown in Fig. 3 was generated by superposing NirF monomers onto the NirS homodimer (PDB: 1nir [5]).

### Size exclusion chromatography with multi angle light scattering (SEC-MALS)

SEC-MALS was used to determine particle sizes in solution. A 10/300 Superdex 200 increase column (GE Healthcare) equilibrated in protein buffer was coupled to an Agilent Technologies 1260 Infinity II HPLC system (Santa Clara, U.S.A.) equipped with a Wyatt Optilab rEX diffraction index detector and a Treos II multi angle laser light scattering detector (Wyatt, Santa Barbara, U.S.A.). Data were processed with ASTRA (7.3.0) utilizing a dn/dc ratio of 0.185 mL/g. Measurements were performed in triplicates.

### Determination of melting temperatures

Protein melting temperatures were determined by difference scanning fluorimetry utilizing the intrinsic fluorescence of tryptophan and tyrosine measured in a Tycho NT6 System (NanoTemper, Munich, Germany). Results are average of three independent measurements with two technical replicates for each measurement.

### Dihydroheme *d*_1_ binding studies

Binding of dd1 to *P. aeruginosa* NirF was assessed with two UV-Vis-based methods. Because spectral features associated with binding were found to be more pronounced at lower pH, the protein was rebuffered to 50 mM BisTris pH 6.5, 500 mM NaCl and 10% (v/v) glycerol using ZeBa spin columns (ThermoFisher Scientific). First, 30 μL of solutions at 40 μM of dd1 and 40 μM NirF were subjected to SEC using the setup described for MALS experiments, albeit with a 5/150 Superdex 75 increase column (GE Healthcare). Elution was followed by measuring the absorption at 395 nm. The area under the curve for co-eluted dd1 was calculated and normalized against the complete signal. Second, remaining samples were diluted to 5 μM and UV/Vis spectra were recorded between 340 and 800 nm using an EvolutionBio 260 UV/Vis spectrophotometer (ThermoFisher Scientific). The spectra shown in Fig. 7 were normalized, averaged and smoothed (Savitzky-Golay algorithm). These experiments were performed three times, including controls without protein or dd1 in the sample.

## Supporting information

Complete supporting materials

## Acknowledgements

We thank the beamline staff at P11 at the PETRAIII synchrotron (Deutsches Elektronensynchrotron DESY, Hamburg, Germany) and at BM29 (European Synchtroton Radiation Facility ESRF, Grenoble, France) for letting us use their facilities. We acknowledge Alejandro Arce Rodriguez and Susanne Häußler for their support in collecting growth curves of the *P. aeruginosa* strains and Sabrina Hafke for support during protein purification. This work was supported by grants from the Deutsche Forschungsgemeinschaft to GL (LA 2412/3-2) and WB (PROCOMPAS graduate school, GRK 2223/1).

## Abbreviations

DDSH: 12,18-didecarboxysiroheme
dd1: dihydroheme *d*_1_
LB: lysogeny broth
MBP: maltose binding Protein
MALS: multi angle light scattering
Pa.: *Paracoccus*
P.: *Pseudomonas*
PEG: polyethylene glycol
PCR: polymerase chain reaction
R_g_: radius of gyration
SAM: S-Adenosyl methionine
SEC: size exclusion chromatography
SAXS: small angle x-ray scattering
SUMO: small ubiquitin-related modifier
tNCS: translational non-crystallographic symmetry

