## Supplementary material for "Crystal structure of NirF: Insights into its role in heme *d*_1_ biosynthesis": Complete supporting materials

**Supplementary Table 1:** Data collection and refinement statistics

| | MALX-E-NirF | NirF | NirF dd1 |
|---|---|---|---|
| **Data collection statistics** | | | |
| Resolution shell type | spherical | ellipsoidal | ellipsoidal |
| Wavelength (Å) | 1.000000 | 1.000000 | 1.736500 |
| Space group | P 1 $2_1$ 1 | P 1 $2_1$ 1 | P 1 $2_1$ 1 |
| Unit cell dimensions | | | |
| a, b, c (Å) | 76.09, 117.60, 103.31 | 101.3, 147.83, 108.72 | 101.20, 149.44, 110.17 |
| α, β, γ (°) | 90, 90.175, 90 | 90, 98.21, 90 | 90, 98.48, 90 |
| Resolution range (Å)[a] | 117.6-4.0 (4.47-4.00) | 107.60-1.56 (1.73-1.56) | 100.09-1.89 (2.14-1.89) |
| Ellipsoidal resolution (Å) (direction)[h] | - | 1.70 (0.930**a***+0.367**c***) 1.56 (**b***) 2.11 (-0.41**a***+0.912**c***) | 2.10 (0.955**a***-0.296**c***) 1.89 (**b***) 2.76 (0.13**a***+0.992**c***) |
| Total no. of reflections[a] | 63534 (17507) | 1073574 (54339) | 918555 (24628) |
| No. of unique reflections[a] | 15238 (4329) | 289596 (14420) | 149789 (7472) |
| Multiplicity[a] | 4.2 (4.0) | 3.7 (3.8) | 6.1 (3.3) |
| Completeness (%) (spherical)[a] | 98.9 (98.9) | 65.2 (69.2) | 58.8 (9.9) |
| Completeness (%) (ellipsoidal)[a,h] | - | 94.7 (69.3) | 93.0 (52.6) |
| $R_{p.i.m}$ (all I+&I-)[a,c] | 0.467 (0.728) | 0.067 (0.394) | 0.076 (0.342) |
| I/σ(I)[a] | 2.9 (1.8) | 6.4 (1.7) | 6.1 (1.9) |
| $CC_{1/2}$[a,c] | 0.255 (0.182) | 0.995 (0.655) | 0.991 (0.742) |
| Largest Patterson peak with length larger than 15 Å (tNCS) | | | |
| Fraction coordinates[i] | - | 0.5, 0.276, 0.5 | 0.5, 0.285, 0.5 |
| Height relative to origin (%)[i] | - | 38.529 | 41.565 |
| **Refinement statistics** | | | |
| No.of refections used in refinement[a] | - | 289582 (2508) | 149796 (623) |
| No.of refelctions used for $R_{free}$[a] | - | 14291 (117) | 7422 (37) |
| $R_{work}$[a,d] | - | 0.1628 (0.2472) | 0.2153 (0.3085) |
| $R_{free}$[a,e] | - | 0.2024 (0.2989) | 0.2576 (0.3155) |
| No.of protein residues | - | 2976 | 2976 |
| No. of non hydrogen atoms | - | 27416 | 25402 |
| protein | - | 23789 | 23477 |
| ligand | - | 58 | 392 |
| solvent | - | 3569 | 1533 |
| Average B-factor (Å$^2$)[g] | - | 20.76 | 22.87 |
| Protein | - | 19.11 | 22.74 |
| Water | - | 41.62 | 21.13 |
| Ligands | - | 31.44 | 25.32 |
| RMSD[g] | - | | |
| Bond length (Å) | - | 0.006 | 0.010 |
| Bond angle (°) | - | 0.87 | 1.09 |

| | | | |
|---|---|---|---|
| Ramachandran plot (%)[g] | - | | |
| Favored regions | - | 98.38 | 97.13 |
| Allowd regions | - | 1.62 | 2.87 |
| outliers | - | 0.00 | 0.00 |
| Rotamer outliers (%) | - | 0.20 | 0.36 |
| Cis Prolines (%) | | 10.13 | 10.13 |
| Clash score[f] | - | 3.31 | 2.65 |
| MolProbity score[f] | - | 1.05 | 1.17 |
| **PDB Code** | | 6TV2 | 6TV9 |

[a]Values in parentheses are for the highest resolution shell.

[b]$R_{\text{p.i.m}} = \Sigma_{hkl}\{1/[N(hkl) - 1]\}^{1/2} \times \Sigma_i|I_i(hkl) - \langle I(hkl)\rangle|/\Sigma_{hkl}\ \Sigma_i\ I_i(hkl)$.

[c]$CC_{1/2} = \Sigma(x - \langle x\rangle)(y - \langle x\rangle)/[\Sigma(x - \langle x\rangle)^2\Sigma(y - \langle y\rangle)^2]^{1/2}$.

[d]$R_{\text{work}} = (\Sigma_{hkl}||F_{\text{obs}}| - k\ |F_{\text{calc}}||)/(\Sigma_{hkl}\ |F_{\text{obs}}|)$.

[e]$R_{free}$ is the same as $R_{\text{work}}$ with 5% of reflections chosen at random and omitted from refinement.

[f]Statistics calculated with the MolProbity web server [1] (http://molprobity.biochem.duke.edu/)

[g]Statistics calculated with Table 1 tool of PHENIX suite [2] |).

[h]Statistics calculated with StaranisoServer [3]

[i]Calculated with Xtriage [4]

**Supplementary Table 2:** Plasmids used in this work

| Plasmid name | Description | Reference |
|---|---|---|
| pUCP20T | *Escherichia-Pseudomonas* shuttle vector | West *et al*. [5] |
| pUCP20T-*nirF* | *nirF* gene cloned pUCP20T | Nicke *et al*. [6] |
| pET-SUMO | pET19m (modified pET19 (Invitrogen)) with SUMO tag cloned in Nde1 restriction site in frame with 6xHis and TEV site | Lab Collection |
| pET-SUMO-*nirF* | *nirF* without export signal from pUCP20T-*nirF* cloned in between Nde1 and BamHI restriction site in frame with SUMO tag | This study |
| pMALX-E | Modified pMAL-c2x (New England Biolabs) containing for crystallization enhanced maltose binding protein | Moon *et al*. [7] |
| pMALX-E-*nirF* | *nirF* without export signal from pUCP20T-nirF cloned in between Nhe1 and HindIII restriction site in frame with MBP | This study |
| pUCP20T-*nirF*$^{R21A}$ | pUCP20T-*nirF* with R21 mutated to alanine | This study |
| pUCP20T-*nirF*$^{H48A}$ | pUCP20T-*nirF* with H48 mutated to alanine | This study |
| pUCP20T-*nirF*$^{S50A}$ | pUCP20T-*nirF* with S50 mutated to alanine | This study |
| pUCP20T-*nirF*$^{R65A}$ | pUCP20T-*nirF* with R65 mutated to alanine | This study |
| pUCP20T-*nirF*$^{D185A}$ | pUCP20T-*nirF* with D185 mutated to alanine | This study |
| pUCP20T-*nirF*$^{Y234A}$ | pUCP20T-*nirF* with Y234 mutated to alanine | This study |
| pUCP20T-*nirF*$^{K235A}$ | pUCP20T-*nirF* with K235 mutated to alanine | This study |
| pUCP20T-*nirF*$^{H238A}$ | pUCP20T-*nirF* with H238 mutated to alanine | This study |
| pUCP20T-*nirF*$^{E240A}$ | pUCP20T-*nirF* with E240 mutated to alanine | This study |
| pUCP20T-*nirF*$^{H325A}$ | pUCP20T-*nirF* with H325 mutated to alanine | This study |
| pUCP20T-*nirF*$^{R340A}$ | pUCP20T-*nirF* with R340 mutated to alanine | This study |
| pUCP20T-*nirF*$^{R372A}$ | pUCP20T-*nirF* with R372 mutated to alanine | This study |
| pET-SUMO-*nirF*$^{R21A}$ | pET-SUMO-*nirF* with R21 mutated to alanine | This study |
| pET-SUMO-*nirF*$^{H48A}$ | pET-SUMO-*nirF* with H48 mutated to alanine | This study |
| pET-SUMO-*nirF*$^{R65A}$ | pET-SUMO-*nirF* with R65 mutated to alanine | This study |
| pET-SUMO-*nirF*$^{D185A}$ | pET-SUMO-*nirF* with D185 mutated to alanine | This study |
| pET-SUMO-*nirF*$^{K235A}$ | pET-SUMO-*nirF* with K235 mutated to alanine | This study |
| pET-SUMO-*nirF*$^{H238A}$ | pET-SUMO-*nirF* with H238 mutated to alanine | This study |
| pET-SUMO-*nirF*$^{R372A}$ | pET-SUMO-*nirF* with R372 mutated to alanine | This study |

**Supplementary Table 3:** Strains used in this work

| Strain name | Description | Reference |
|---|---|---|
| E.coli BL21 | *Escherichia coli* host used for recombinant protein expression | |
| E.coli XL1-blue | *Escherichia coli* host used for cloning | |
| PA01 | *Pseudomonas aeruginosa* PA01 | Lab Collection |
| PA01 RM361 | *Pseudomonas aeruginosa* PA01 *nirN::tet* | Kawasaki *et al.* [8] |
| PA01 RM301 | *Pseudomonas aeruginosa* PA01 *nirF::tet* | Kawasaki *et al.* [9] |
| BL21-Sumo-*nirF* | *E.coli* BL21 with pET-SUMO-*nirF* | This study |
| BL21-Sumo-$nirF^{R21A}$ | *E.coli* BL21 with pET-SUMO-$nirF^{R21A}$ | This study |
| BL21-Sumo-$nirF^{H48A}$ | *E.coli* BL21 with pET-SUMO-$nirF^{H48A}$ | This study |
| BL21-Sumo-$nirF^{R65A}$ | *E.coli* BL21 with pET-SUMO-$nirF^{R65A}$ | This study |
| BL21-Sumo-$nirF^{D185A}$ | *E.coli* BL21 with pET-SUMO-$nirF^{D185A}$ | This study |
| BL21-Sumo-$nirF^{K235A}$ | *E.coli* BL21 with pET-SUMO-$nirF^{K235A}$ | This study |
| BL21-Sumo-$nirF^{H238A}$ | *E.coli* BL21 with pET-SUMO-$nirF^{H238A}$ | This study |
| BL21-Sumo-$nirF^{R372A}$ | *E.coli* BL21 with pET-SUMO-$nirF^{R372A}$ | This study |
| PA-wt | PA01 with PUCP20T | This study |
| PA-Δ*nirF* | PA01 RM301 with pUCP20T | This study |
| PA-*nirF* | PA01 RM301 with pUCP20T-*nirF* | This study |
| PA-R21A | PA01 RM301 with pUCP20T-$nirF^{R21A}$ | This study |
| PA-H48A | PA01 RM301 with pUCP20T-$nirF^{H48A}$ | This study |
| PA-S50A | PA01 RM301 with pUCP20T-$nirF^{S50A}$ | This study |
| PA-R65A | PA01 RM301 with pUCP20T-$nirF^{R65A}$ | This study |
| PA-D185A | PA01 RM301 with pUCP20T-$nirF^{D185A}$ | This study |
| PA-Y234A | PA01 RM301 with pUCP20T-$nirF^{Y234A}$ | This study |
| PA-K235A | PA01 RM301 with pUCP20T-$nirF^{K235A}$ | This study |
| PA-H238A | PA01 RM301 with pUCP20T-$nirF^{H238A}$ | This study |
| PA-E240A | PA01 RM301 with pUCP20T-$nirF^{E240A}$ | This study |
| PA-H325A | PA01 RM301 with pUCP20T-$nirF^{H325A}$ | This study |
| PA-R340A | PA01 RM301 with pUCP20T-$nirF^{R340A}$ | This study |
| PA-R372A | PA01 RM301 with pUCP20T-$nirF^{R372A}$ | This study |

**Supplementary Table 4:** Primers used in this work; the parts of a primer that match the sequence of NirF are in bold letters. Mismatching base pairs of primers used in mutagenesis are highlighted in bold red.

| Primer name | Sequence (5'→3') |
|---|---|
| nirF_pET-SUMO_fw | CGCGAACAGATCGGTGGTCATATG**ATGAGCCAGCAGCCGC** |
| nirF_pET-SUMO_rv | GCTTTGTTAGCAGCCGGATCCTA**GAGTCCGATGTGCTGGGCG** |
| nirF_pMALX-E_fw | CAGACTAATGCGGCCGCAGCTAGC**ATGAGCCAGCAGCCGC** |
| nirF_pMALX-E_rv | CGACGGCCAGTGCCAAGCTT**AGAGTCCGATGTGCTGGGCG** |
| nirF_R21A_rv | CCGTCGGCG**GC**TTCGATCAGCACGCCGAG |
| nirF_R21A_fw | TCGGCGTGCTGATCGAA**GC**CGCCGACGGCAG |
| nirF_H48A_rv | CACCAGGGAGGCG**GC**GGACAGGTCGCCG |
| nirF_H48A_fw | CGGCGACCTGTCC**GC**CGCCTCCCTGGTG |
| nirF_S50A_rv | GAGAACACCAGGG**C**GGCGTGGGACAGG |
| nirF_S50A_fw | CCTGTCCCACGCC**G**CCCTGGTGTTCTC |
| nirF_R65A_rv | GCCGCCGTCG**GC**ACCGAATACGTAGGCGTAG |
| nirF_R65A_fw | TACGCCTACGTATTCGGT**GC**CGACGGCGGC |
| nirF_D185A_rv | GGCTGATGAGGGCG**G**CGTAGGGTTGCTTG |
| nirF_D185A_fw | CAAGCAACCCTACG**C**CGCCCTCATCAGCC |
| nirF_Y234A_rv | GGTGCGGCATCTTG**GC**CACCGGCAGCTTGC |
| nirF_Y234A_fw | GCAAGCTGCCGGTG**GC**CAAGATGCCGCACC |
| nirF_K235A_rv | CCAGGTGCGGCATC**GC**GTACACCGGCAGCTTG |
| nirF_K235A_fw | AGCTGCCGGTGTAC**GC**GATGCCGCACCTGG |
| nirF_H238A_rv | GTCCAGCCCTCCAGG**GC**CGGCATCTTGTACAC |
| nirF_H238A_fw | GTGTACAAGATGCCG**GC**CCTGGAGGGCTGGAC |
| nirF_E240A_rv | TGGTCCAGCCC**G**CCAGGTGCGGC |
| nirF_E240A_fw | GCCGCACCTGG**C**GGGCTGGACCATC |
| nirF_H325A_rv | CGCTGAATTCCATG**GC**CAGTACGCCGGGGC |
| nirF_H325A_fw | GCCCCGGCGTACTG**GC**CATGGAATTCAGCG |
| nirF_R340A_rv | CTGGTCGGCGTCG**GC**CACCGAGATCCAG |
| nirF_R340A_fw | CTGGATCTCGGTG**GC**CGACGCCGACCAG |
| nirF_R372A_rv | CGATGTGCTGGGCG**GC**GTGGCTGAAGAAGATG |
| nirF_R372A_fw | TCTTCTTCAGCCAC**GC**CGCCCAGCACATCG |

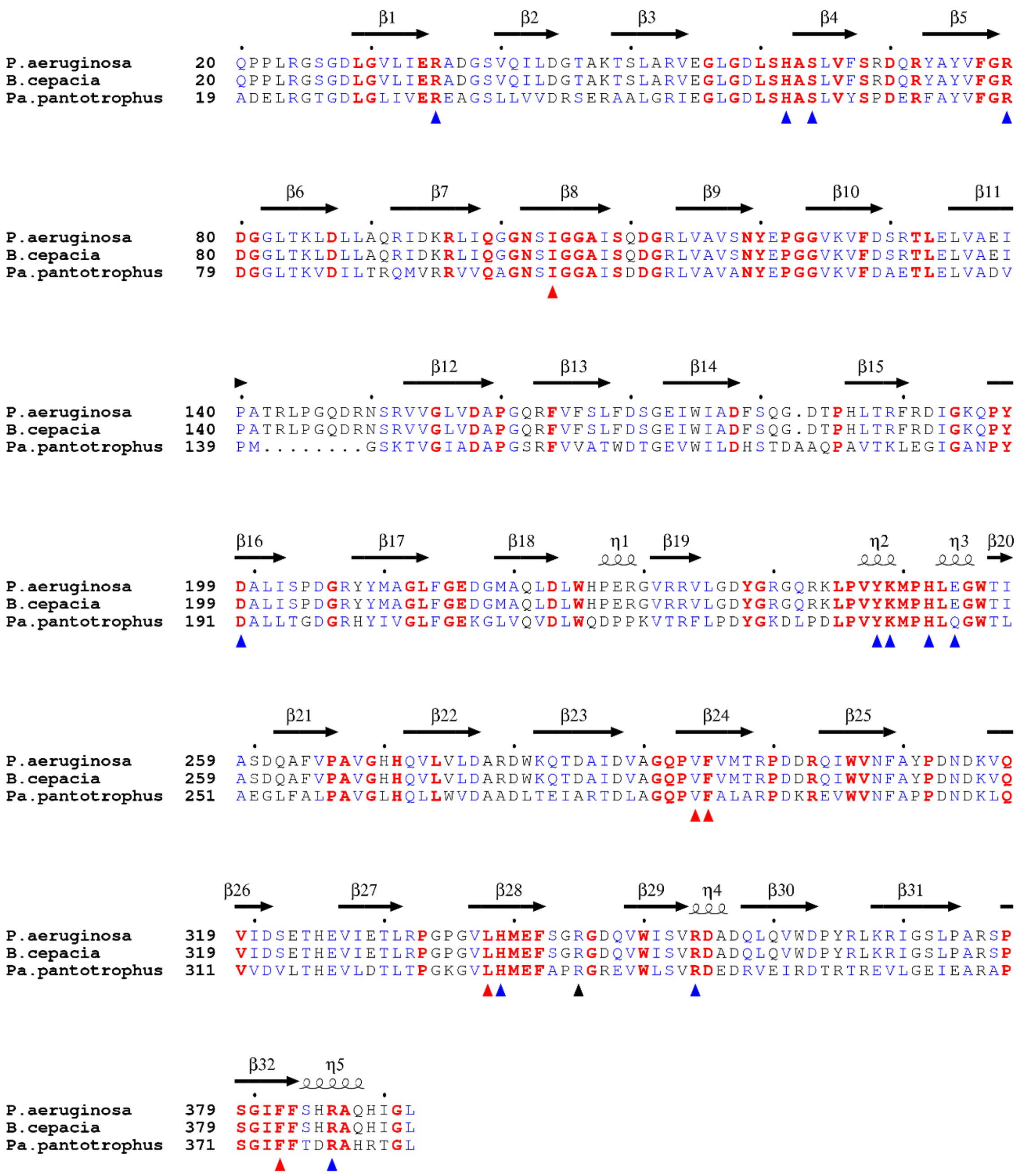


**Supplementary Figure 1:** Sequences of NirF from 21 organisms (seven of each σ-,β- or γ-proteobacteria) aligned with ClustalΩ [10] and depicted with ESPript3.0 [11]. Only one representative sequence for each division is depicted. Numbering of sequences is based on complete sequences including the signal peptide. Red bold letters mark identity in all sequences and blue a 70% conservation as calculated with the Risler matrix. Secondary structure elements of NirF without bound dd1 are depicted on top of the sequences. Blue arrows on the bottom mark residues mutated in this study, red highlights hydrophobic residues interacting with dd1 and black arrows indicates R331, which is part of the dimer interface. The complete sequence alignment is provided in a separate file.

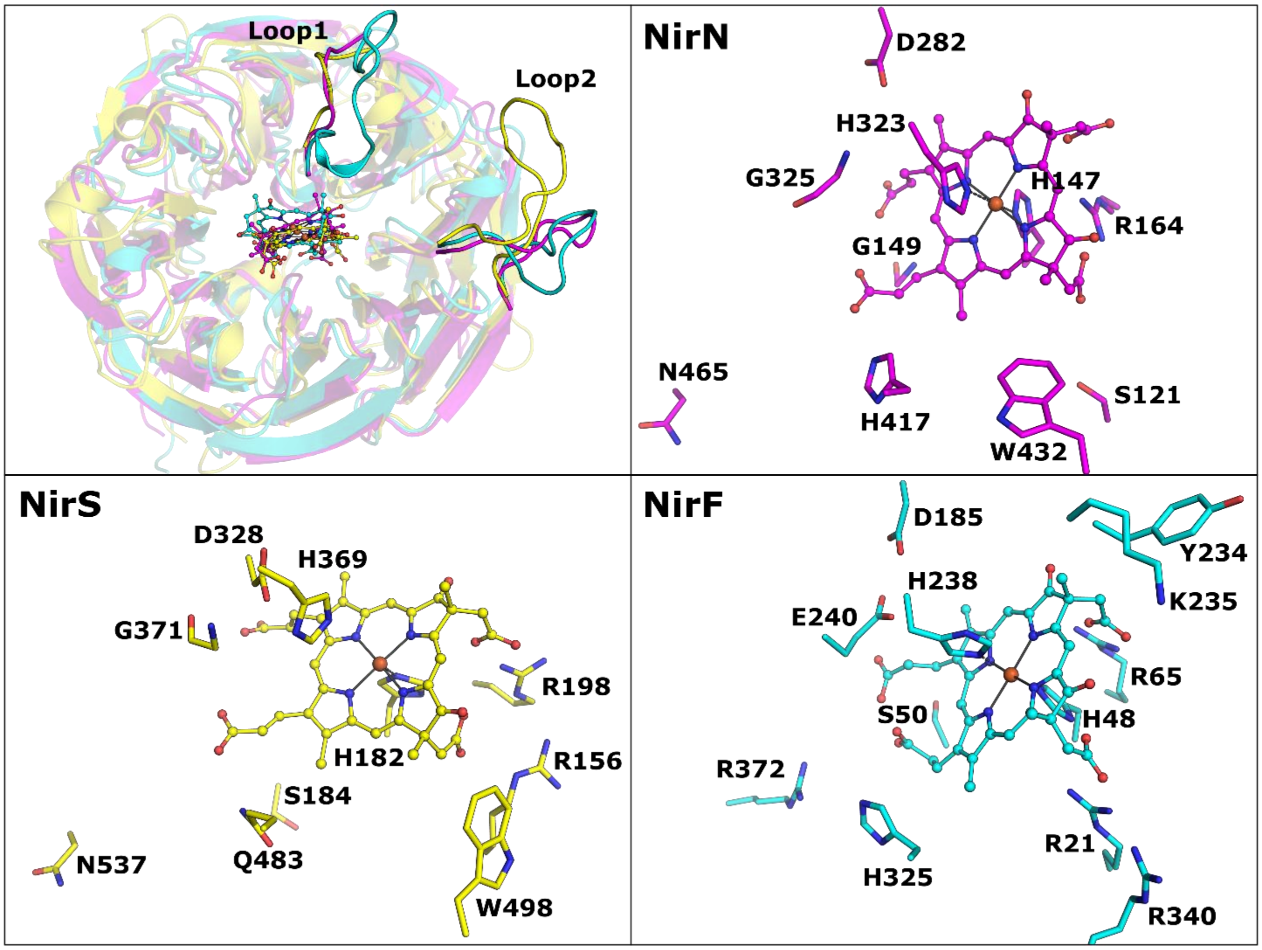


**Supplementary Figure 2:** Comparison of d1-domain of NirF (cyan), NirN (magenta) and NirS (yellow) after superposition with PyMol.

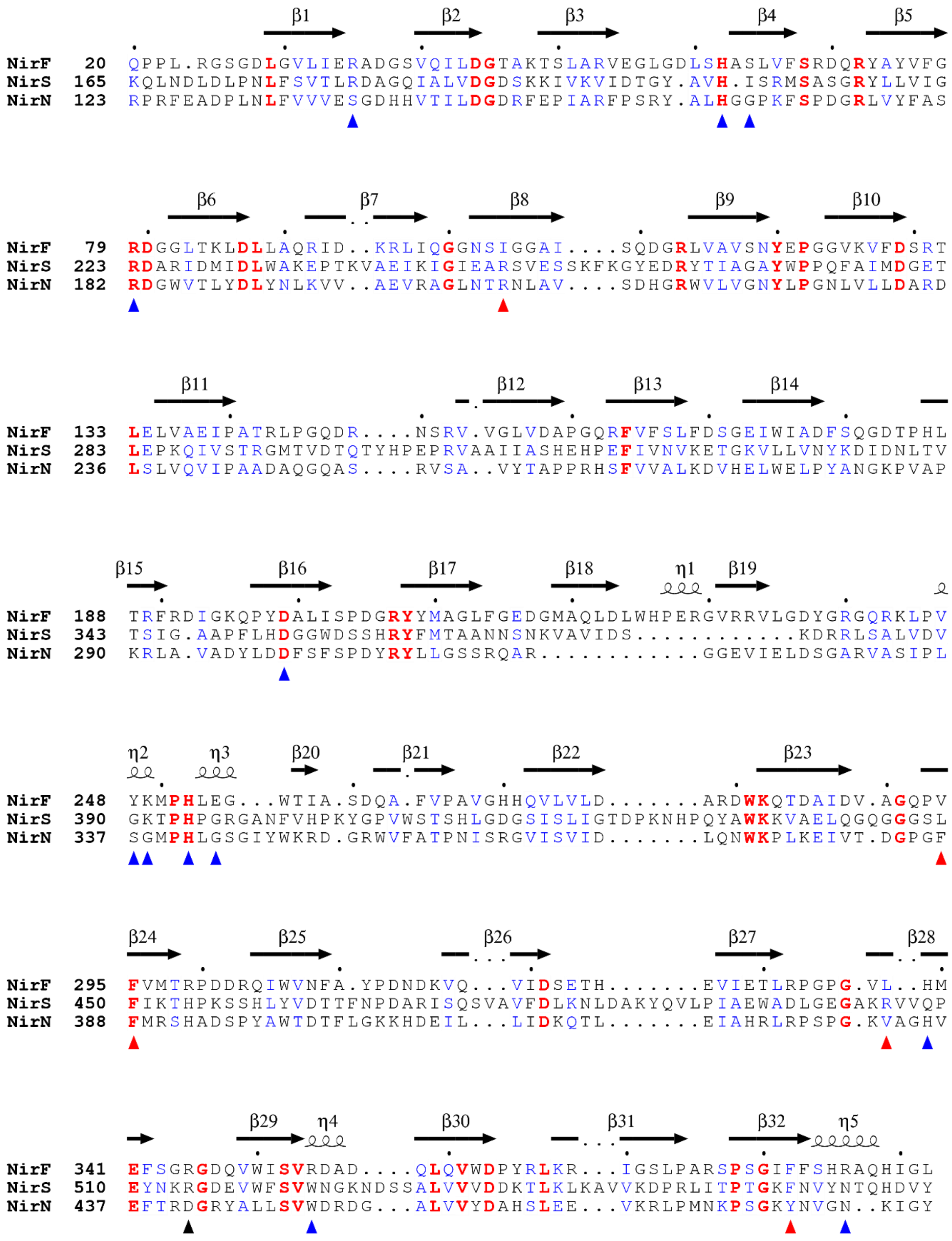


**Supplementary Figure 3:** Sequence alignment of the d1-domain of NirF, NirS and NirN from *P. aeruginosa*. Depiction according to Supplementary Figure 1.
